# Modulating the proliferative and cytotoxic properties of human TIL by a synthetic immune niche of immobilized CCL21 and ICAM1

**DOI:** 10.1101/2022.07.07.499105

**Authors:** Sharon Yunger, Benjamin Geiger, Nir Friedman, Michal J Besser, Shimrit Adutler-Lieber

**Affiliations:** Ella Lemelbaum Institute for Immuno-Oncology, Sheba Medical Center, Ramat Gan, Israel; Department of Immunology, the Weizmann Institute of Science, Rehovot, Israel; Department of Clinical Microbiology and Immunology, Sackler School of Medicine, Tel Aviv University, Tel Aviv, Israel

**Keywords:** T-cells, Tumor Infiltrating Lymphocytes (TIL), cancer immunotherapy, Synthetic Immune Niche (SIN)

## Abstract

The major challenge in developing an effective adoptive cancer immunotherapy, is the *ex-vivo* generation of tumor-reactive cells, in sufficient numbers and with enhanced cytotoxic potential. It was recently demonstrated, that culturing of activated murine CD8+ T-cells on a “Synthetic Immune Niche” (SIN), consisting of immobilized CCL21 and ICAM-1, enhances T-cell expansion, increases cytotoxicity against cultured cancer cells and suppresses tumor growth *in vivo* [1, 2]. In the study reported here, we have tested the effect of the CCL21+ICAM1 SIN, on the expansion and cytotoxic phenotype of Tumor Infiltrating Lymphocytes (TIL), following activation with immobilized anti-CD3/CD28 stimulation, or commercial activation beads. The majority of TIL tested, displayed higher expansion when cultured on the coated SIN compared to cells incubated on uncoated substrate. Comparable enhancement of TIL proliferation was obtained by the CCL21+ICAM1 SIN, in a clinical setting that includes a 14-day rapid expansion procedure (REP), initiated with feeder cells, anti CD3 and IL-2. Co-incubation of post-REP TIL with matching target cancerous cells, demonstrated increased IFNγ secretion beyond baseline in most of the TILs and a significant increase in granzyme B levels following activation on SIN. The SIN did not significantly alter the relative frequency of CD8/CD4 populations, as well as the expression of CD28, CD25 and several exhaustion markers. These results demonstrate the potential capacity of the CCL21+ICAM1 SIN to reinforce TIL-based immunotherapy.

## Introduction

Adoptive cancer immunotherapy, employing ex vivo expanded genetically-modified or natural autologous T-cells, is currently one of the most promising approaches used for the treatment of cancer patients. [7-3] Adoptive therapy approaches in cancer patients are mainly based on the administration of either Chimeric Antigen Receptor (CAR)-expressing T-cells [8, 9] or Tumor Infiltrating Lymphocytes (TILs) [10-12].

CAR T-cells are polyclonal T-cells, usually isolated from the patient’s peripheral blood, that are genetically engineered to express a CAR with the specificity of a monoclonal antibody, directed against a known tumor-associated antigen (TAA), along with a co-stimulatory signaling capacity. [13, 14]

Conversely, TIL-based therapy uses the natural infiltration of T-cells into tumors, as indication for their tumor antigen recognition capacity, and their immunotherapeutic potential. TIL, that originally display insufficient proliferative and cytotoxic functionality under the suppressive conditions within the tumor, are isolated from the resected tumor tissue, and cultured *ex-vivo* under supportive restimulating conditions. Briefly, the *ex-vivo* production process of TIL consists of two major stages [15-17]: During the first stage, referred to as pre-Rapid Expansion Procedure (pre-REP), TIL cultures are being established, by the resection of a tumor biopsy, outgrowth of the tumor resident lymphocytes and their expansion, induced by interleukin-2 (IL-2) added to the culture medium. In the next stage - REP, the established TIL culture is massively expanded to treatment levels by addition of soluble anti-CD3 antibody, IL-2, and irradiated feeder cells. By the end of the REP, TILs are administered back to the lympho-depleted cancer patient.

While there are unique advantages to TIL therapy over other adoptive cellular therapies, including its broad TCR diversity, relative safety and effective tumor homing, there are still major challenges to consider for its wide application, including varying clinical response rates in treating different patients and different types of tumors, variations in the REP expansion yields, unpredicted expansion of T-cell clones with different therapeutic capacities [18] and the risk that prolonged TIL expansion may result in impaired functionality, such as T-cell exhaustion or anergy [19-21].

Towards reinforcing adoptive cancer immunotherapy, a ‘Synthetic Immune Niche’ (SIN) strategy was recently developed by Adutler-Lieber et.al., based on the stimulation of T-cells by surfaces, coated with the chemokine C-C motif Ligand 21 (CCL21) and the Intercellular Adhesion Molecule 1 (ICAM-1) [1, 2].

CCL21, secreted by lymphatic stroma and endothelial cells [22] induces several processes critical to immune responses, including co-localization and recruitment of T-cells and dendritic cells (DCs); [23, 24] facilitating cell migration; [25, 26] priming T-cells for synapse formation; [27] and co-stimulation of naïve T-cell expansion and Th1 polarization. [28-30] ICAM1 plays a key role in the formation of immune synapses and promotion of T cell activation, through binding to its integrin ligand, lymphocyte function-associated 1 (LFA1).[31, 32]

These factors are expected to act synergistically, as CCL21 increases LFA1 responsiveness to ICAM1, and mediates the arrest of motile lymphocytes on ICAM1-expressing DCs and endothelial cells, and their clustering with other T-cells. [22, 33] Indeed, the developed SIN, consisting of a combination of immobilized CCL21 and ICAM-1, was shown to increase the expansion of activated murine CD4+ [2] and CD8+ [1] T-cells. Furthermore, culturing of OVA-specific CD8+ T-cells on this SIN elevates the cellular levels of granzyme B expressed by these cells and increases their efficiency in killing ovalbumin-expressing cultured cancer cells, as well as their tumor suppressive activity, in vivo. [1]

In the present study we apply, for the first time, the CCL21+ICAM1 based SIN technology, on human TILs, derived from melanoma patients. We show here that SIN-treatment of anti CD3/CD28 activated cells display higher expansion rates than cells incubated on uncoated substrate. Comparable enhancement of TIL proliferation was also obtained when the SIN treatment was included in the rapid expansion process. Furthermore, testing of selected, SIN-treated, post-REP TIL cultures, demonstrated increased IFNγ secretion and granzyme B expression levels, suggesting that the CCL21+ICAM1 SIN can reinforce the TIL-based immunotherapeutic system.

## Materials and methods

### Generation and expansion of TIL cultures

TIL samples were obtained from patients enrolled to an open-label phase II ACT trial for patients with metastatic melanoma at the Sheba Medical Center (NCT00287131).

The establishment of TIL cultures was performed as previously described [15-17]. In short, fragmentation, enzymatic digestion and cell remnants technique were used to isolate TIL from surgically resected metastatic lesions. During the first week, non-adherent TIL were transferred to a new 24-well plate, and cultured separately from the adherent melanoma cells. TILs were cultured in complete medium (CM) containing 10% human AB serum (Valley Biomedical, Winchester, VA or Gemini Bio, West Sacramento, CA), 2mM L-glutamine (Biological Industries, Israel), Pen/Strep (Biological Industries, Israel) and 3,000 IU/ml IL-2 (Chiron Novartis, New Jersey, USA) in RPMI 1640 medium (Gibco, Thermo Fisher Scientific, Waltham, MA). Cells were split or medium was added every 2 to 3 days, to maintain a cell concentration of 0.5–2.0 × 10^6^ cells/ml. TIL cultures were established within 2 to 4 weeks.

### Stimulation of TIL with anti CD3/CD28 antibodies

TILs were stimulated for 7 days with anti-biotin MACSiBead Particles loaded with biotinylated CD3/CD28 antibodies (Miltenyi Biotec, Bergisch Gladbach, Germany). The preparation of the conjugated beads was carried out according to the manufacture’s recommendation (activation/expansion kit, Miltenyi Biotec). Antibody-loaded anti-biotin MACSiBead™ were added to the TIL at a cell/bead ratio of 1:1. In addition TIL were stimulated for 7 days with 2 μg/ml plate-bound anti-CD3 (clone OKT-3; Miltenyi Biotech) and 2 μg/ml anti-CD28 (clone: 15E8, Miltenyi Biotech).

### Surface coating and functionalization with CCL21 and ICAM-1

Substrate functionalization was performed by overnight incubation with 5 µg/ml CCL21 and 50 µg/ml ICAM1 (R&D Systems, Minneapolis, MN, USA), in PBS.

### Rapid Expansion Procedure (REP) with CCL21 and ICAM-1 surface coating

The REP was initiated by stimulating 0.01-0.03 ×10^6^ TIL with 30 ng/ml anti-CD3 antibodies (MACS GMP CD3 pure, clone OKT-3; Miltenyi Biotech, Germany), 3,000 IU/ml IL-2, and irradiated peripheral blood mononuclear cells from three non-related donors as feeder cells (50 Gy, TIL to feeder cells = 1:100) in 50% CM, 50% AIM-V medium (Invitrogen, Thermo Fisher Scientific, Waltham, MA). REP was performed in 24-well plates and CCL21+ICAM1 coating was prepared one day prior to the initiation or cell splitting (days −1, 6 and 10). TILs were cultured for 14 days and split on day 7 and 11 to maintain a cell concentration of 0.3-2.0×10^6^/well.

### Antibodies and flow cytometry

The following antibodies were used for flow cytometry: Pacific blue-CD3 (clone SK7, BioLegend, San Diego, CA, United States), PECy7 or FITC-CD8 (clone HIT8a, BioLegend), FITC-PD-1 (clone EH12.2H7, BioLegend), FITC-TIM-3 (clone F38-2E2, BioLegend), FITC-LAG-3 (clone 11C3C65, BioLegend) APC-CD25 (clone BC96, BioLegend), APC-CD28 (clone CD28.2, BioLegend), APC-vio770-CD45RA (clone HI100, BioLegend) and PerCP-CCR7 (clone G043H7, BioLegend). TIL were washed and re-suspended in cell-staining buffer (BioLegend). Cells were incubated for 30 min with the antibodies on ice, washed in buffer and measured using MACSQuant flow cytometer (Miltenyi Biotech).

For localization and quantification of granzyme B, cells were fixed (BLG420801, Biolegend), permeabilized (BLG421002, Biolegend), and stained with granzyme B-specific antibodies (BLG515406, BioLegend). For carboxy fluorescein succinimidyl ester (CFSE) cell proliferation assays, TIL were stained before seeding at day 11 of the REP with 5 μM CFSE (Thermo Fisher Scientific) for 20 min at 37°C, according to the manufacturer’s instructions. Four days later cells were taken for flow cytometry analysis and their CFSE fluorescence intensity was determined. Samples were analyzed using FlowJo software (FlowJo LLC, Ashland, OR).

### IFNγ ELISA

1×10^5^ TIL were co-cultured with autologous melanoma cell lines in 96-well plates overnight at 1:1 effector-to-target ratio. Supernatant was collected and IFNγ levels were determined by ELISA (BioLegend) according to the manufacture’s protocol.

### Statistics

Significance of variation between groups was evaluated using a non-parametric two-tailed Student’s t-test. Test for differences between proportions was performed using two-sided Fisher’s exact test with p ≤ 0.05 values considered as significant.

## Results

### Impact of CCL21+ICAM1 coated surface on TIL proliferation, following CD3/CD28 stimulation

To evaluate the effect of a CCL21+ICAM1 coated surface on the proliferative capacity of TIL, the cells were seeded in a 24-well plate, coated with CCL21+ICAM1 (“coated-TIL”) or a non-coated plate (“uncoated-TIL”) and activated by CD3/CD28 beads stimulation. Six TIL cultures were tested. TIL cultures were derived from five patients with metastatic melanoma (Pt. 014, 124, 132, 145 and 151) and two, independently-established, TIL cultures from the same patient (Pt. 014/F3 and 014/F4). Seven days following the initial stimulation, five, out of the six TIL cultures, showed higher fold expansion values following exposure to CCL21+ICAM1 **(Figure 1A and 1D)**. In these five cultures, the expansion of coated-TIL was, on average, 2.5 ± 1.2-fold higher (range 1.6-4.6, p=0.026) after 7 days, achieving an 8.4 ± 7.5-fold expansion for coated-TIL compared with 4.5 ± 8.5-fold for TIL cultured on uncoated surfaces.

**Figure 1.**
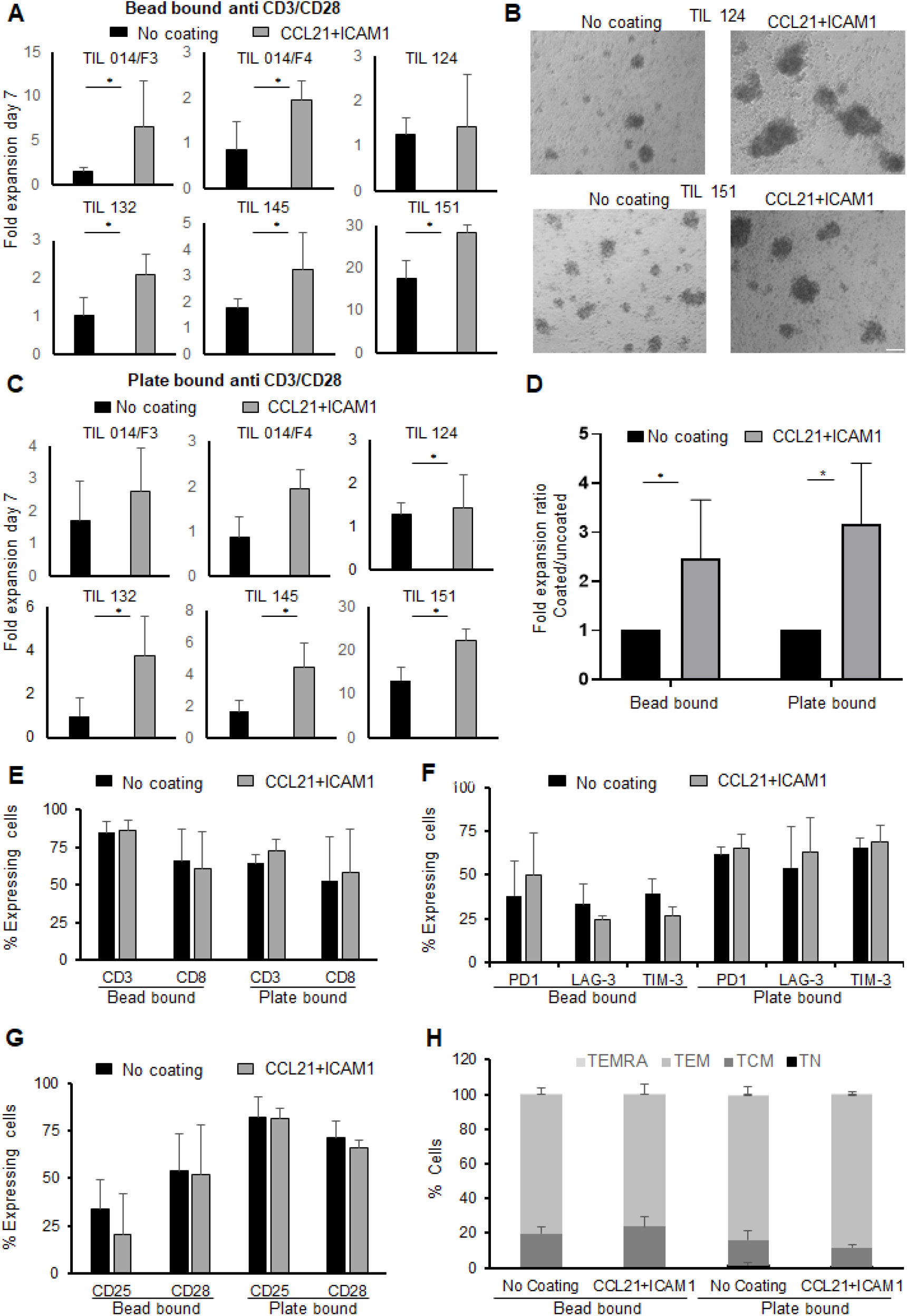
Fold expansion and phenotype analysis following anti-CD3/CD28 stimulation of TIL with or without exposure to CCL21+ICAM1 coated surface. Fold expansion of TIL cultures stimulated by **(A)** anti-CD3/CD28 activation beads or **(B)** Morphology following 7-day plate-bound CD3/CD28 activation (scale bar = 100 µm). **(C)** Plate-bound anti-CD3/CD28 antibodies for 7 days. Graph bars shows comparison between TIL cultured on uncoated vs. CCL21+ICAM1-coated surfaces.**(D)** Average fold expansion of TIL cultures, stimulated with either activation beads or plate-bound anti CD3/CD28 antibodies after normalization (no-coated TIL = 1, n = 6). **(E)** Percentage of CD3 and CD8 cells **(F-G)** Distribution of surface markers: co-inhibitory: PD1, LAG-3, TIM-3, activation marker: CD25 and co-stimulatory CD28. **(H)** TIL differentiation status: TN (naïve), CD3+CD45RA+CCR7+; TCM (central memory), CD3+CD45RA−CCR7+; TEM (effector memory), CD3+CD45RA−CCR7−; TEMRA (effector), CD3+CD45RA+CCR7.

Moreover, microscopy-based monitoring demonstrated that SIN-stimulated TIL tend to form larger cell aggregates than those incubated on non-coated substrates **(Figure 1B)**. In addition, when stimulating TIL cultures with plate-bound anti CD3 and anti CD28 antibodies (instead of activation beads), four out of six TIL cultures exposed to CCL21+ICAM1 demonstrated 3.2 ± 1.2-fold higher proliferation (p = 0.012) over TIL cultured on uncoated dishes (fold expansion of 8.1 ± 9.4 versus 4.0 ± 6.1, range 1.69-4.49, **Figure 1C-D)**. By the end of the expansion, most of the cells (65-85%) were CD3+ T cells. Expansion of anti CD3 and anti CD28 stimulated TIL on CCL21+ICAM1 coated plates had no apparent impact on the CD4/CD8 subpopulation distribution **(Figure 1E**, p = 0.815) or the co-inhibitory molecules PD-1 (p = .800), LAG-3 (p = .511) and TIM-3 (p = .680), on the expression of the co-stimulatory CD28 (p = 0.475), the activation marker CD25 (p = .942) and the differentiation status defined by the expression of CD45RA in combination with CCR7 (p ≥ 0.05) **(Figure 1F-H)**. In addition, there was no significant difference in the expression of sub-population markers within the CD4 or CD8 populations (p-values ≥ 0.05) **(Suppl. Table 1 and 2)**. The gating strategy and representative flow cytometric plots are shown in **Supplementary Figure 1**.

**Table 1.**
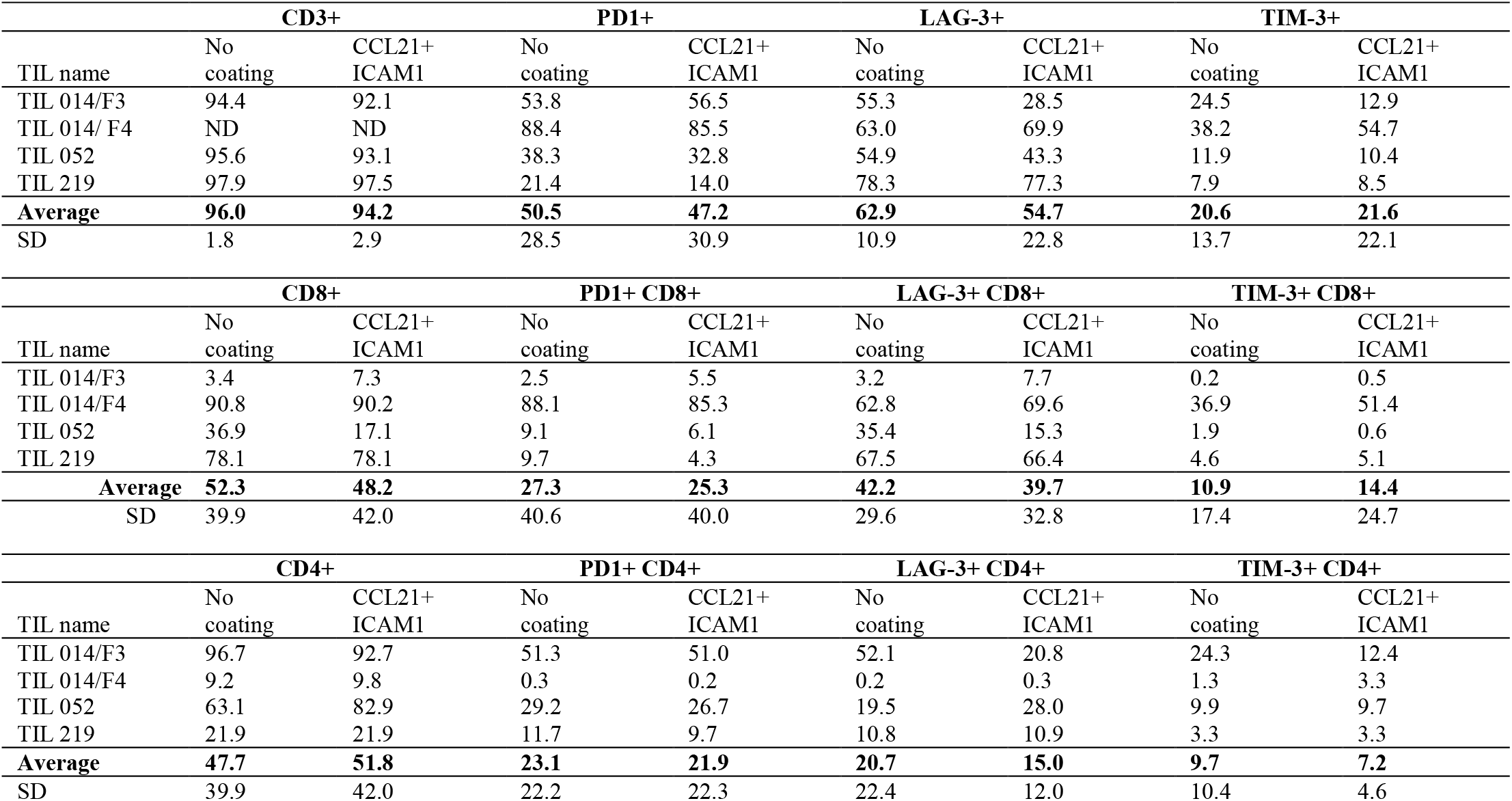
Phenotype analysis of post-REP TIL, cultured on CCL21+ICAM1-coated or uncoated surfaces.

### CCL21+ICAM1 coated surfaces increased TIL proliferation during the rapid expansion procedure (REP)

In the clinical setting, TIL are expanded rapidly to large cell numbers by adding 40-50 Gy irradiated PBMC feeder cells, soluble anti-CD3 antibody and IL-2 to the cell culture. On day 14 of the ex-vivo expansion, TIL are harvested and prepared for the i.v. infusion. To test whether CCL21+ICAM1-coated surfaces improve the manufacturing process, small-scale REPs were initiated, closely resembling the clinical setting, using GMP-compliant reagents. Six TIL cultures obtained from five melanoma patients (Pt. 014, 052, 219, 031 and 174) were expanded according to the REP protocol for 14 days. As shown in **Figure 2A**, expansion was significantly increased on CCL21+ICAM1-coated surfaces by 2.3 ± 0.7-fold (p=.002) in 5 out of 6 TIL cultures.

**Figure 2.**
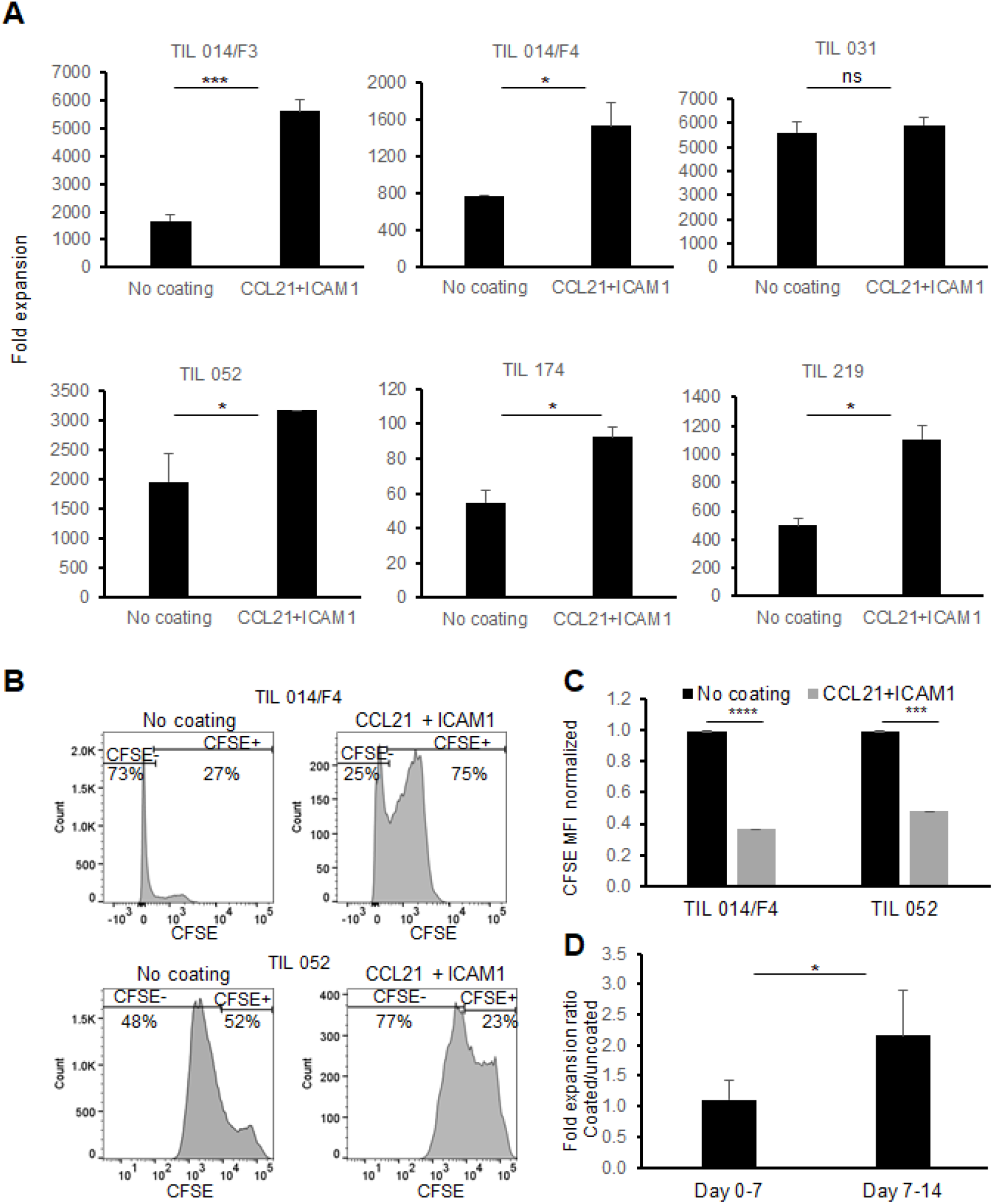
Comparison of TIL during REP with/without CCL21+ICAM1 surface coating. **(A)** Fold expansion on day 14. **(B)** Histogram plots of CFSE staining **(C)** Bar graph shows mean fluorescence intensity of CFSE. TIL cultured on CCL21+ICAM1 coated surfaces normalized to the uncoated TIL (= 1). **(D)** The average frequency of fold expansions at day 0-7 and day 7-14 with or without CCL21+ICAM1 coated surface (n = 5).

To confirm that CCL21+ICAM1 coating contributes to the enhanced expansion by increasing the proliferation rate, TIL were stained with carboxyfluorescein succinimidyl ester (CFSE) dye on day 11 of the REP, and analyzed on day 14. As shown in **Figure 2B-C**, a decrease in CFSE fluorescence intensity was observed in the SIN-stimulated TIL cultures indicating that the increased expansion is attributable to elevated cell proliferation.

It was further shown that the major contribution of the CCL21+ICAM1 coated surface to the increased proliferation occurred during the second week of the REP (fold increase day 0-7 = 1.11 ± 0.32; fold increase day 7-14 = 2.15 ± 0.74; p = .020) **(Figure 2D)**. The impact between day 7 to 11 and day 11 to 14 day was similar (fold increase day 7-11 = 1.61 ± 0.59; fold increase day 11-14: 1.39 ± 0.37; p=0.511).

### Increased expansion of TIL on CCL21+ICAM1-coated surface had no impact on TIL phenotype

To investigate the impact of CCL21+ICAM1-coated surfaces on the TIL phenotype, post-REP TIL (n=7) were analyzed for CD4 and CD8 expression. The average frequency of CD4 and CD8 TIL populations was similar to that of cells cultured on uncoated surfaces (%CD4 cells on coated surfaces= 45.77 ± 34.26%, compared to 43.97 ± 32.24%, p = .658 on uncoated surfaces; %CD8 on coated surfaces = 54.23 ± 34.26%, %CD8 on uncoated surfaces, 56.03 ± 32.24%, p = .927). The CD8 percentage of six TIL cultures prior REP and following REP with or without exposure to CCL21+ICAM1 is shown in **Figure 3A**.

**Figure 3.**
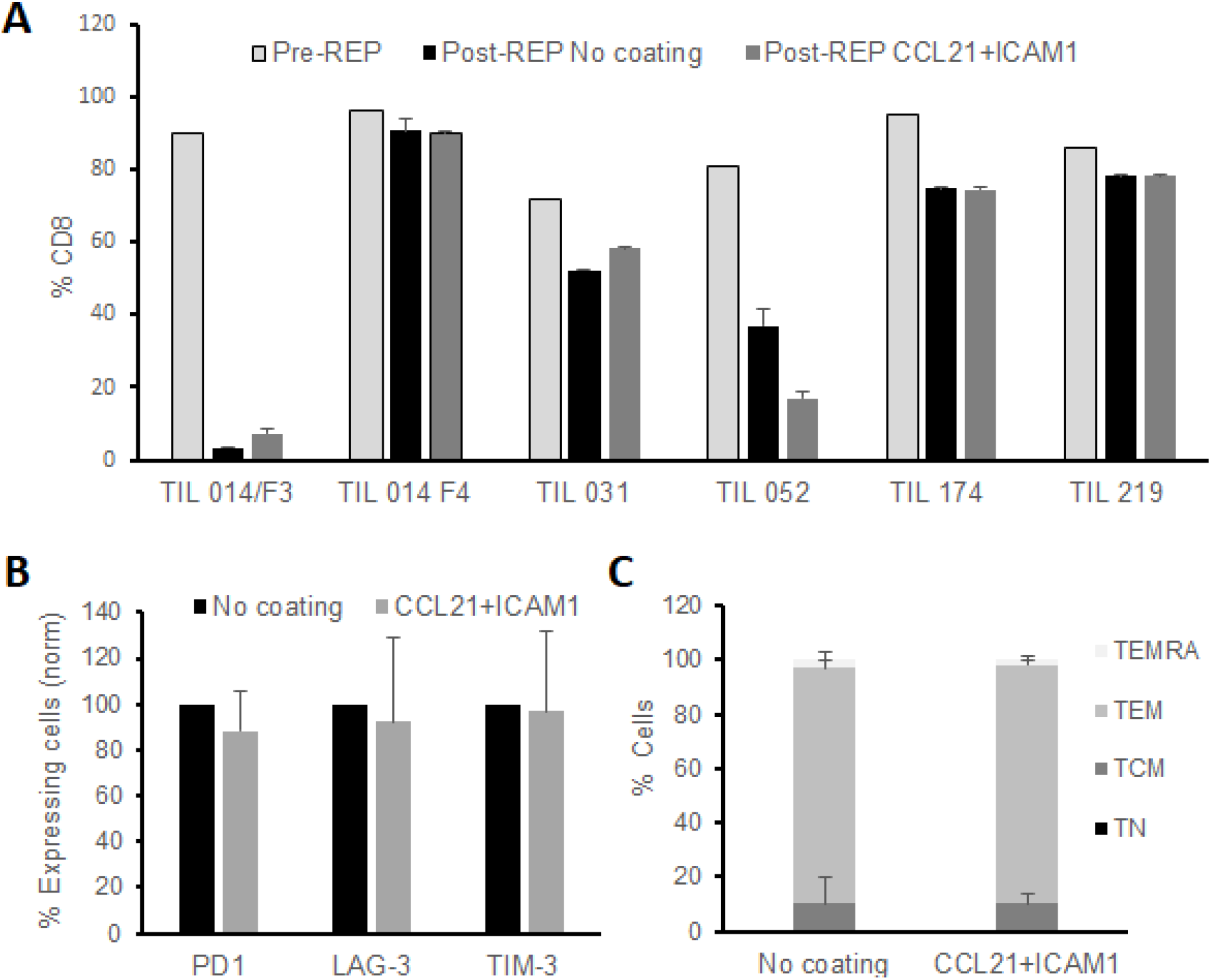
Phenotype analysis of post-REP TIL. **(A)** Frequency of CD8 for both CCL21+ICAM1 coated and uncoated pre and post-REP TIL cultures. **(B)** TIL were characterized for expression of inhibitory molecules PD-1, LAG-3, TIM-3 (n = 4). The expression values of coated surface TIL were normalized to uncoated surface TIL (100%) **(C)** Distribution of differentiation subsets. TN (naive), CD3+CD45RA+CCR7+; TCM (central memory), CD3+CD45RA−CCR7+; TEM (effector memory), CD3+CD45RA−CCR7−; TEMRA (effector), CD3+CD45RA+CCR7−. (n=3).

Four post-REP TIL cultures were further characterized for the expression of the inhibitory molecules PD-1, LAG-3 and TIM-3 on all CD3 T cells or on CD4 / CD8 T cell subpopulations. As shown in **Table 1**, expression of the inhibitory molecules was different in individual TIL cultures, but the difference between cells growing with or without the SIN was insignificant (p-values ≥ .210) **(Figure 3B)**.

The differentiation status of three post-REP TIL cultures was defined based on the expression of CD45RA, in combination with CCR7. The majority of post-REP TIL, independently of whether they were cultured on a CCL21+ICAM1 surface or not, were CD45RA-CCR7-, effector memory T cells (TEM) (coated surfaces, TEM = 88.07 ± 3.36%; uncoated, TEM = 86.73 ± 9.57%; p = .83) **(Figure 3C)**.

### CCL21+ICAM1 coating increased the anti-tumor reactivity of TIL

To determine whether the CCL21+ICAM1-coated surfaces enhance anti-tumor reactivity, IFNγ secretion and granzyme B expression were determined after a co-culture of post-REP TIL cultures with or without autologous tumor target cells.

Interestingly, the basal IFNγ levels of TIL (without target cells) was lower in cells incubated on coated surfaces compared to those cultured on uncoated surfaces **(Figure 4A)**, yet two of four post-REP TIL cultures demonstrated increased IFNγ secretion beyond baseline following exposure to the CCL21+ICAM1-coated surface (**Figure 4B**). Furthermore, SIN-treated TIL 014/F3, co-cultured with autologous tumor cells displayed higher levels of secreted IFNγ (measured by ELISA), as well as increased levels of granzyme B, compared to cells co-cultured with autologous tumor cells on uncoated plates. Notably, in the absence of autologous target cells, the levels of IFNγ, were very low, and the SIN effect on granzyme B levels was limited (**Supplementary Figure S2**).

**Figure 4.**
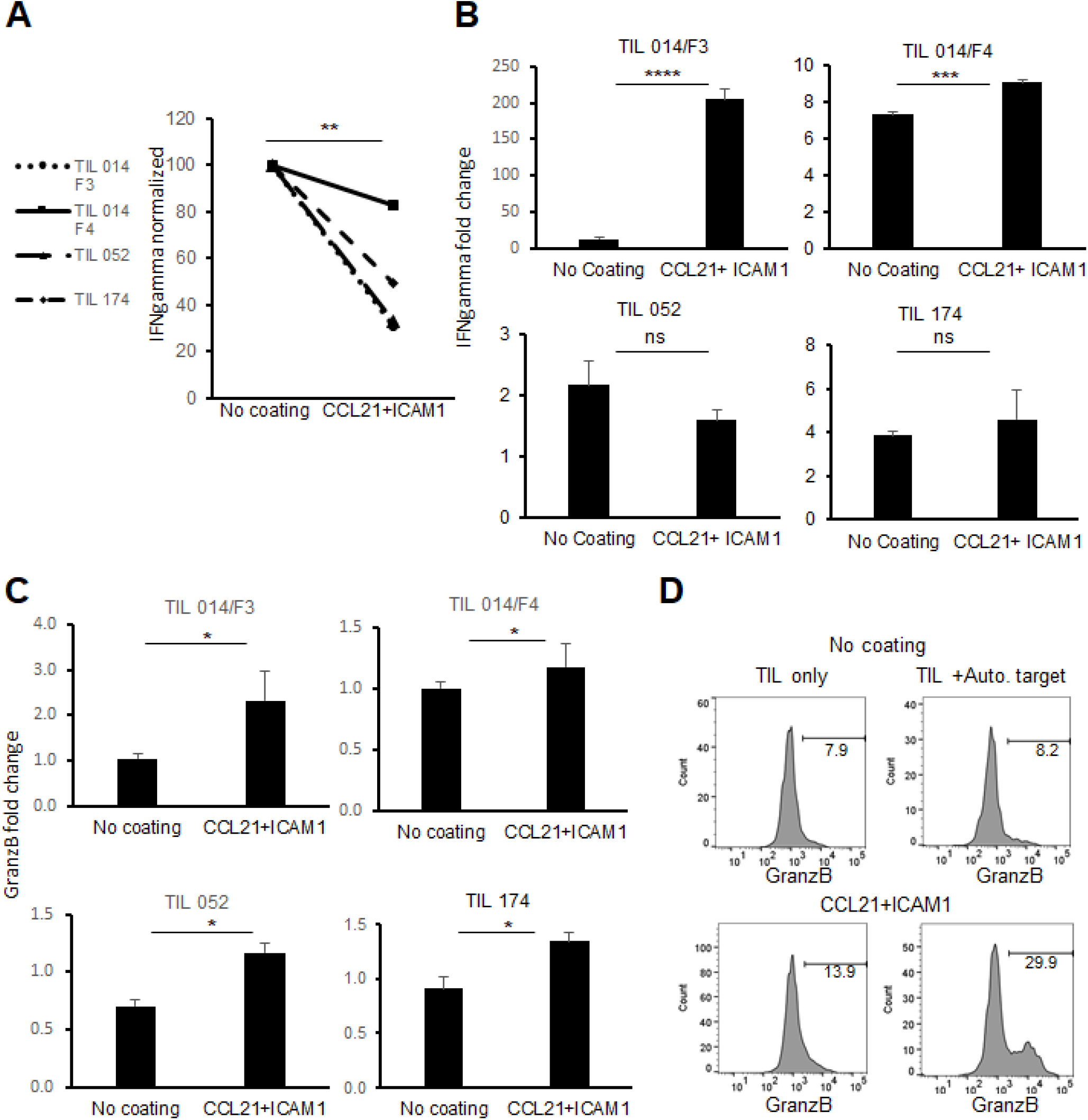
Anti-tumor reactivity and cytotoxicity of TIL cultured on CCL21+ICAM1-coated surfaces. **(A)** Connecting line histogram demonstrated different baseline of IFN-γ secretion from CCL21+ICAM1 coated TIL or uncoated surfaces. **(B)** Fold change of IFN-γ following co-cultures of TIL with autologous tumor versus TIL only. In each experiment, IFN-γ concentration after co-culture was divided by the “TIL only” concentration and expressed as fold change. The bar plots show the mean fold change ± standard deviation (SD) of three independent experiments. **(C)** Fold change of MFI for granzyme B in CD8+ cells co-cultured with tumor cells. MFI values after co-culture were divided by the “TIL only” values, and are presented as the mean ± standard deviation (SD) obtained in three independent experiments. (**D**) Representative FACS Histogram plots for granzyme B.

In addition, granzyme B expression was significantly increased in all tested post REP TIL cultures (n=4) which were expanded on coated surfaces (p ≤ 0.05) **(Figure 4C)**. Representative flow cytometric histograms are shown in **Figure 4D**.

## Discussion

The primary motivation underlying the development an ex-vivo “synthetic immune niche”, that might improve the performance of therapeutic T-cells, is based on current experiments with diverse adoptive immunotherapies, whose effectiveness varies greatly between patients and tumor types [34]. In our earlier studies we have shown, that stimulation of murine CD4+ [1] or CD8+ T-cells [2] by a synthetic immune niche, consisting of immobilized ICAM-1 and CCL21, enhances the proliferation of both cell types, and increases the cytotoxic efficacy of CD8+ cells, both *ex-vivo* and *in-vivo*. These encouraging findings indicated that specific environmental stimulation can alter T-cell proliferation and activity, yet the experimental settings of these studies [35] (murine OT-1 and OT-2 systems), were very different from current clinical settings in humans. The primary goal of the current study was to determine the capacity of human CCL21+ICAM1 SIN to exert similar effects in a clinically relevant system.

The system we chose, namely TIL therapy, has earned great success in treating solid tumors [36], particularly metastatic melanoma, [37-41] and does not necessitate the prediction of an actual TAA, or the characteristics of its binding receptor. TIL cultures are naturally heterogeneous, potentially targeting a variety of tumor antigens, a feature that is particularly relevant to melanoma, which displays a high mutational load [42].

Moreover, the indication that the CCL21+ICAM1 SIN enhances expansion of T-cells, render the SIN approach highly suitable for TIL therapy, in which high numbers of cells are a favorable prognostic indicators for successful therapy in different tumor types, [43-45] and specifically in foretelling the success of checkpoint therapies. [46]. The additional feature of this SIN, based on the results obtained using the murine system, namely, potentiation of the cytotoxic machinery by increasing granzyme B levels [1], is also highly relevant to TIL therapy, in which failed therapies are often attributed to exhaustion and anergy [21, 47].

The results presented herein strongly confirm that using human SIN components, CCL21+ICAM1, to stimulate patient-derived TILs, enhances TIL expansion, when tested in a clinical setting that includes a 14-day rapid expansion procedure, in addition to the standard presence of feeder cells, anti CD3 and IL-2. Importantly, the increased expansion did not impact the differentiation status of TIL or the expression of inhibitory molecules such as TIM-3, LAG-3 and PD-1. Moreover, post-REP TIL demonstrated increased IFNγ secretion beyond baseline in 50% of the cultures and a significant increase in granzyme B levels in all tested TIL (n=4).

Given the results presented here, we propose that the CCL21+ICAM1 synthetic niche, which proved to enhance expansion in the murine OVA systems, and elevate the cytotoxic capacity of murine CD8 cells, is capable to reinforce human TIL, derived from melanoma patients. The mechanism underlying the effect of the CCL21+ICAM1 SIN, is still unclear, yet the results presented here suggest that incorporation of SIN stimulation into the TIL expansion process might significantly improve the therapeutic performance of these cells.

## Acknowledgements

This study was supported by grants from the Israel Science Foundation (program: Israel Precision Medicine Partnership (IPMP) and the Volkswagen Foundation, both to BG and by the generous support of the Lemelbaum family to MJB.

## Figures Legends

**Supplementary Figure 1:**
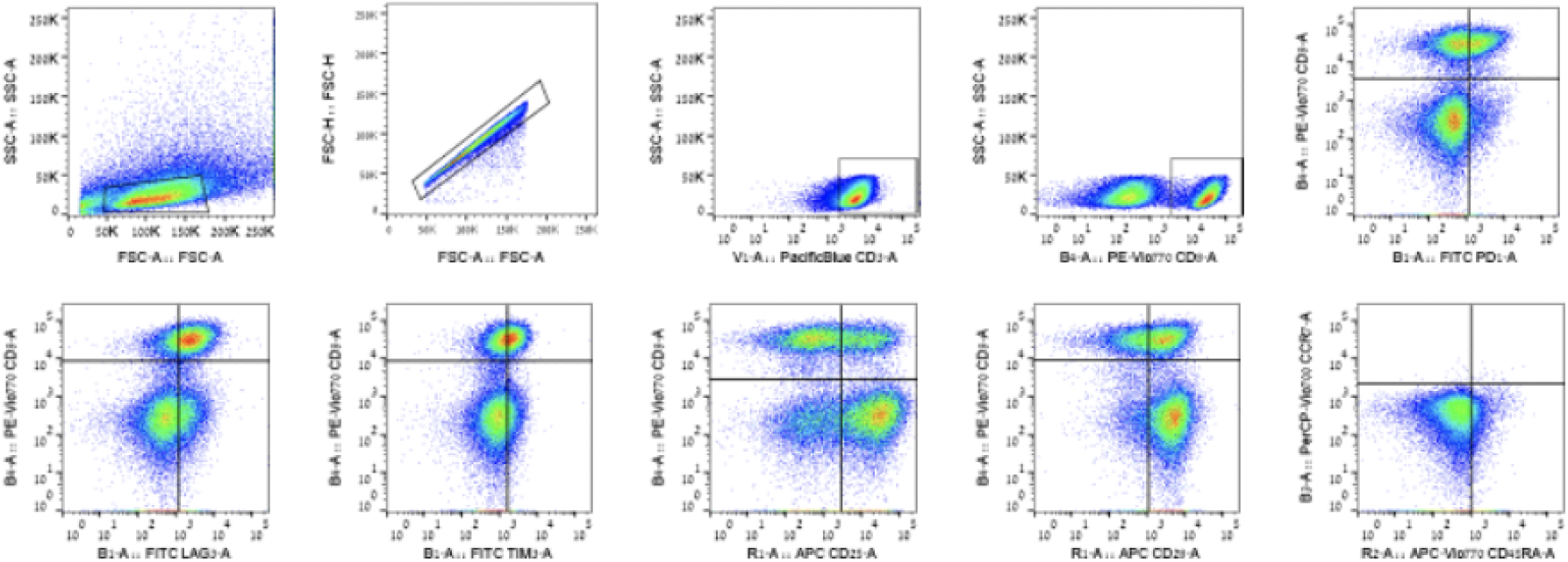
Phenotype analysis of post-REP TIL : Expression of co-inhibitory, co-stimulatory, and differentiation markers. An example of a gating strategy used to analyze the following surface markers: CD3, CD8, PD1, LAG3, TIM3, CD28, CD25 and CD45RAin combination with CCR7. Cells were gated on viable and singlet T cells.

**Supplementary Figure 2.**
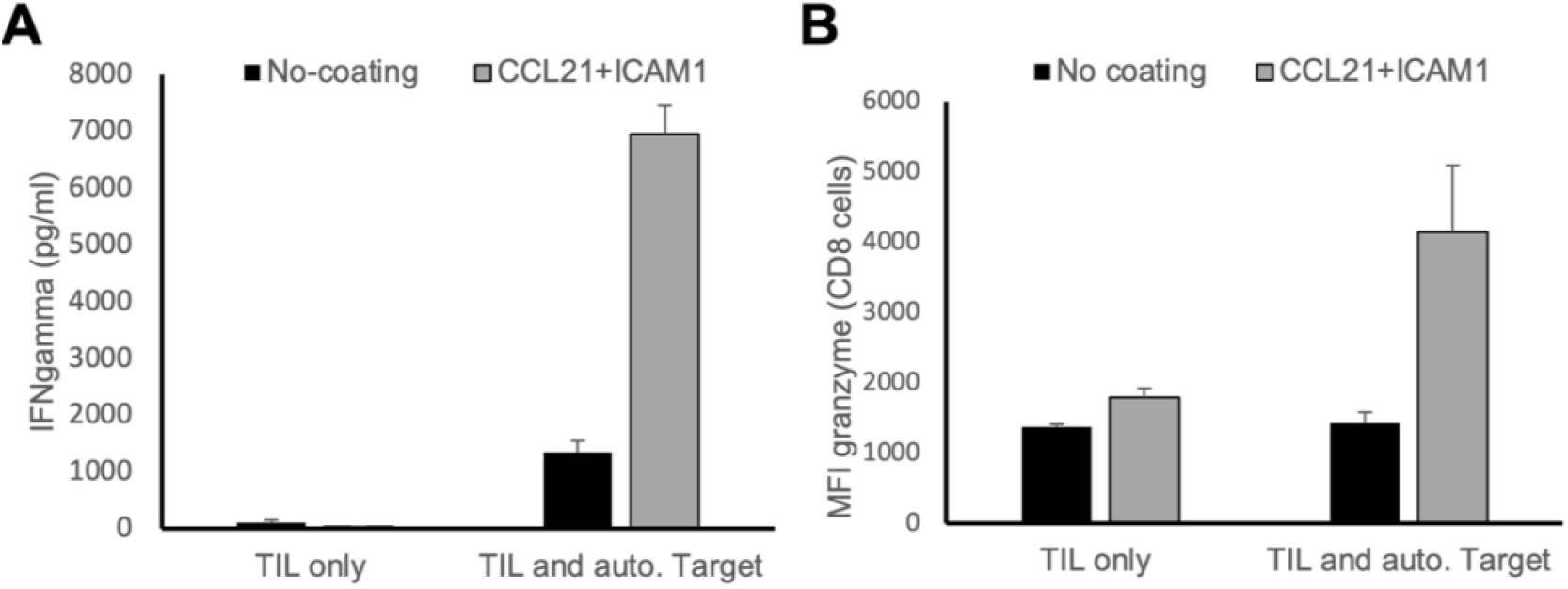
Significant anti-tumor reactivity and cytotoxicity of TIL 014 cultured on CCL21+ICAM1-coated surfaces. (A-B) TIL 014-F3 were co cultures with autologous tumor. Summarizing graphs showing: **(A)** IFNγ secretion (pg/ml) measured by ELISA. **(B)** MFI of granzyme B. Error bars represent the standard deviation of triplicate repetitions

**Supplementary Table 1.**
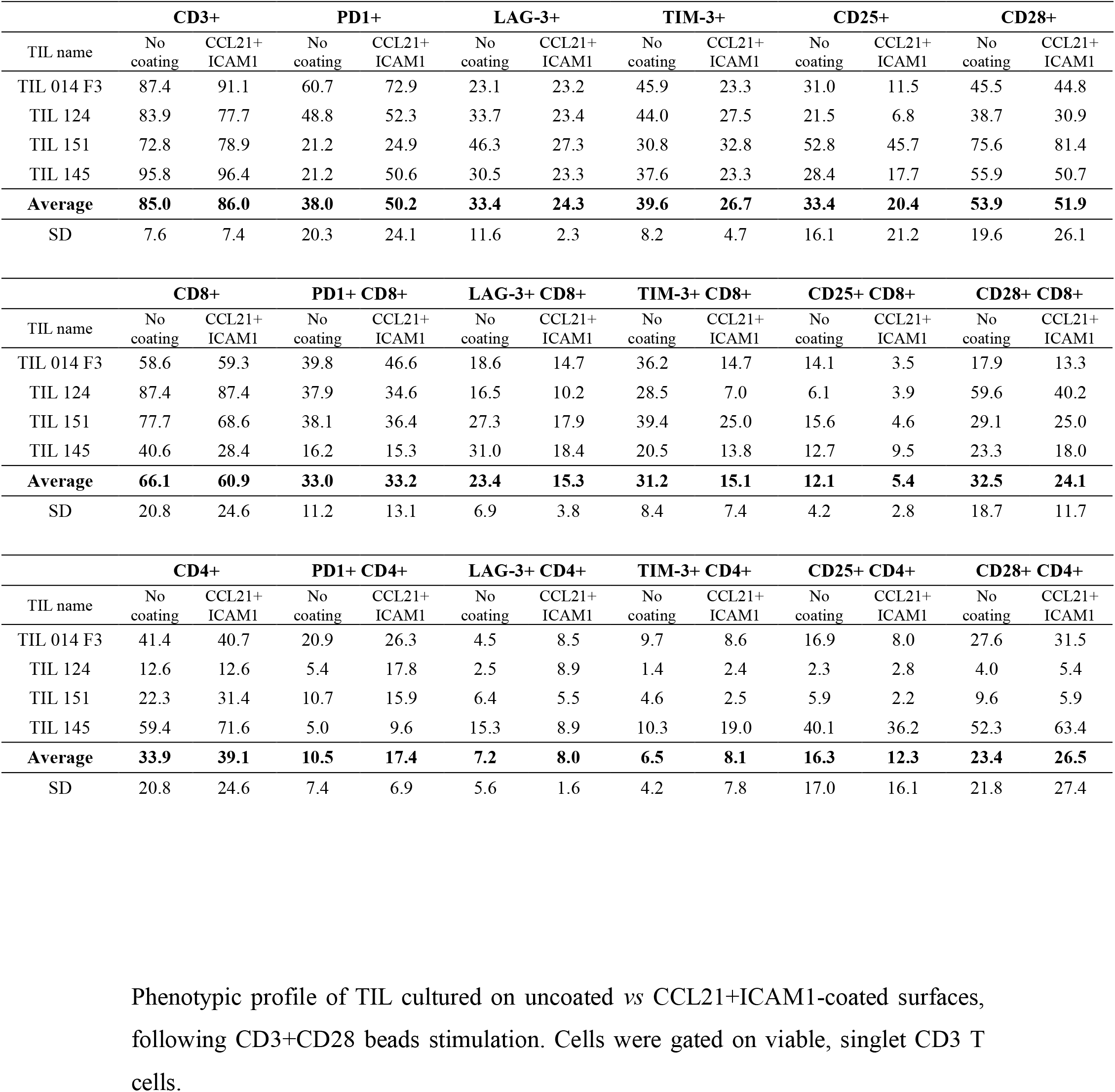

**Supplementary Table 2.**

|  | **CD3+** |  | **PD1+** |  | **LAG-3+** |  | **TIM-3+** |  | **CD25+** |  | **CD28+** |  |
| --- | --- | --- | --- | --- | --- | --- | --- | --- | --- | --- | --- | --- |
| TIL name | No coating | CCL21+ ICAM1 | No coating | CCL21+ ICAM1 | No coating | CCL21+ ICAM1 | No coating | CCL21+ ICAM1 | No coating | CCL21+ ICAM1 | No coating | CCL21+ ICAM1 |
| TIL 014 F3 | 51.4 | 76.6 | 70.7 | 81.7 | 25.9 | 41.3 | 75.2 | 79.4 | 88.2 | 84.4 | 81.6 | 75.7 |
| TIL 124 | 59.9 | 65.7 | 67.5 | 65.8 | 65.6 | 67.7 | 65.1 | 62.1 | 77.8 | 75.0 | 72.4 | 69.8 |
| TIL 151 | 61.6 | 78.4 | 75.4 | 76.3 | 67.7 | 80.9 | 65.9 | 78.0 | 66.4 | 75.7 | 63.1 | 67.6 |
| TIL 145 | 85.7 | 71.9 | 33.4 | 37.7 | 56.5 | 61.3 | 56.2 | 55.2 | 95.8 | 90.8 | 67.1 | 50.7 |
| **Average** | **64.7** | **73.2** | **61.8** | **65.4** | **53.9** | **62.8** | **65.6** | **68.7** | **82.0** | **81.5** | **71.0** | **66.0** |
| SD | 5.5 | 6.9 | 4.0 | 8.1 | 23.6 | 20.2 | 5.6 | 9.6 | 10.9 | 5.2 | 9.3 | 4.2 |

|  | **CD8+** |  | **PD1+ CD8+** |  | **LAG-3+ CD8+** |  | **TIM-3+ CD8+** |  | **CD25+ CD8+** |  | **CD28+ CD8+** |  |
| --- | --- | --- | --- | --- | --- | --- | --- | --- | --- | --- | --- | --- |
| TIL name | No coating | CCL21+ ICAM1 | No coating | CCL21+ ICAM1 | No coating | CCL21+ ICAM1 | No coating | CCL21+ ICAM1 | No coating | CCL21+ ICAM1 | No coating | CCL21+ ICAM1 |
| TIL 014 F3 | 19.0 | 33.4 | 13.7 | 26.8 | 10.9 | 23.9 | 13.6 | 27.0 | 9.9 | 17.7 | 11.7 | 18.7 |
| TIL 124 | 84.6 | 85.1 | 51.9 | 55.5 | 59.4 | 62.6 | 58.0 | 58.8 | 66.2 | 63.2 | 68.9 | 64.1 |
| TIL 151 | 68.6 | 81.4 | 52.1 | 64.4 | 52.1 | 71.6 | 52.7 | 71.3 | 33.6 | 55.7 | 40.5 | 55.8 |
| TIL 145 | 40.6 | 33.0 | 10.8 | 20.5 | 17.1 | 29.1 | 15.7 | 24.7 | 19.1 | 28.8 | 10.7 | 10.5 |
| **Average** | **53.2** | **58.2** | **32.1** | **41.8** | **34.9** | **46.8** | **35.0** | **45.5** | **32.2** | **41.4** | **33.0** | **37.3** |
| SD | 29.2 | 28.9 | 23.0 | 21.4 | 24.4 | 23.8 | 23.6 | 23.2 | 24.7 | 21.6 | 27.7 | 26.6 |

|  | **CD4+** |  | **PD1+ CD4+** |  | **LAG-3+ CD4+** |  | **TIM-3+ CD4+** |  | **CD25+ CD4+** |  | **CD28+ CD4+** |  |
| --- | --- | --- | --- | --- | --- | --- | --- | --- | --- | --- | --- | --- |
| TIL name | No coating | CCL21+ ICAM1 | No coating | CCL21+ ICAM1 | No coating | CCL21+ ICAM1 | No coating | CCL21+ ICAM1 | No coating | CCL21+ ICAM1 | No coating | CCL21+ ICAM1 |
| TIL 014 F3 | 81.0 | 66.6 | 57.0 | 54.9 | 15.0 | 17.4 | 61.6 | 52.4 | 78.3 | 66.7 | 69.9 | 57.0 |
| TIL 124 | 15.4 | 14.9 | 15.6 | 10.3 | 6.2 | 5.1 | 7.1 | 3.3 | 11.6 | 11.8 | 3.5 | 5.7 |
| TIL 151 | 31.4 | 18.6 | 23.3 | 11.9 | 15.6 | 9.3 | 13.2 | 6.7 | 32.8 | 20.0 | 22.6 | 11.8 |
| TIL 145 | 59.4 | 67.0 | 22.6 | 17.2 | 39.4 | 32.2 | 40.5 | 30.5 | 76.7 | 62.0 | 56.4 | 40.2 |
| **Average** | **46.8** | **41.8** | **29.6** | **23.6** | **19.1** | **16.0** | **30.6** | **23.2** | **49.9** | **40.1** | **38.1** | **28.7** |
| SD | 29.2 | 28.9 | 18.6 | 21.1 | 14.2 | 11.9 | 25.3 | 22.9 | 33.1 | 28.2 | 30.5 | 24.1 |

Phenotypic profile of TIL, cultured on uncoated vs CCL21+ICAM1-coated surfaces, following activation with plate-bound CD3+CD28. Cells were gated on viable, singlet CD3 T cells.

